# The Hindu Kush, not the Indus Valley, divides amphibian biogeographic realms

**DOI:** 10.64898/2026.03.29.715076

**Authors:** Daniel Jablonski, Muhammadullah Rasekh, Arifullah Zia, Mohammad Arif Irfan, Faizurrahman Khalili, Nazar Mohammad Wahaj, Borhanudin Noori, Abdul Rahman Osmani, Naveed Sahil Stanekzai, Rafaqat Masroor, Christophe Dufresnes

## Abstract

The boundary between the Palearctic and Oriental biogeographic realms is one of the oldest unresolved problems in Eurasian biogeography. In Central and South Asia, this transition is remarkable and has often been linked to the Indus River Valley, yet the role of adjacent mountain systems and arid landscapes has remained largely untested because critical regions, especially Afghanistan, have been chronically under-sampled due to difficult, long-term socio-political situation. Here we use a unique amphibian material from Afghanistan and adjacent northwestern Pakistan to test whether the Hindu Kush marks the effective boundary between these two realms. Combining molecular barcoding across all seven amphibian genera inhabiting the region with updated distribution records, we find a strikingly consistent lineage turnover centered on the eastern Hindu Kush and adjacent arid systems. Palearctic taxa reach their southeastern limits along the massif, whereas Oriental taxa extend only to its southern foothills. By contrast, Oriental lineages cross the Indus River without detectable phylogeographic break, indicating that the river valley does not function as a major biogeographic boundary for the taxa examined. The Hindu Kush also harbors endemic relict taxa of both Palearctic and Oriental affinities, revealing a dual role as both refugium and dispersal barrier. Thus, our results identify the Hindu Kush as the de facto boundary between the Palearctic and Oriental realms in this part of Eurasia and show that under-sampled arid-montane regions can disproportionately improve global biogeographic frameworks.

## Introduction

How biodiversity is structured across Eurasia remains one of the oldest yet most debated questions in biogeography. The continent encompasses some of the most pronounced environmental gradients on Earth, from humid tropics to cold deserts and high mountain or Arctic systems, traditionally divided into two major biogeographic realms, the Palearctic and the Oriental. Yet the transition between these realms remains poorly resolved. Since Wallace (1), the region between Central and South Asia has been regarded as a broad transitional zone (2-5), with subsequent hypotheses placing the boundary in markedly different locations around the Indus River Valley depending on the taxonomic group considered (6,7). These discrepancies highlight the complexity of biogeographic responses in this region – shaped by its exceptional topographic and environmental heterogeneity – but also the scarcity of empirical data from adjacent, alternative candidate areas.

One of this unexplored area is the Hindu Kush mountain range and its arid surroundings, which forms a formidable natural barrier spanning remote and extreme environments across Afghanistan and northwestern Pakistan. Decades of restricted access, political unrest, and logistical constraints have hindered biodiversity research in these countries (8). Biological material suitable for phylogeographic analyses have thus remained exceedingly rare, and as a result, the role of the Hindu Kush as a biogeographic transition remains largely unresolved.

Amphibians provide a suitable group to address these questions, given their terrestrial habits, relatively low dispersal, as well as hydric and thermal ecological constraints, which makes them especially responsive to geoclimatic barriers. Biogeographic patterns in amphibians also challenge long-standing assumptions: several species of Oriental origin seem to cross the Indus River Valley and reach the southeastern foothills of the Hindu Kush and adjacent desert margins (9,10), whereas Palearctic lineages seldom extend beyond northern slopes of the range.

Accordingly, although the Indus River can reach widths of ∼1.5 km, its highly dynamic channel morphology and frequent flooding likely make it permeable to dispersal for water-associated organisms. In contrast, the high elevations of the Hindu Kush and the hyper-aridity of surrounding deserts may constitute a more persistent and substantive filter.

To test the role of the Hindu Kush as the de facto boundary between the Palearctic and Oriental realms, we here gathered and analyzed unique amphibian material collected during recent field surveys across the Hindu Kush, with emphasis on historically inaccessible regions. Using molecular barcoding, we identify and map lineages across all seven amphibian genera inhabiting the area, and combined with updated occurrence records for Afghanistan, we assess the biogeographic boundaries between the Palearctic and Oriental realms in this underexplored part of Eurasia.

## Results

Amphibian diversity and distribution patterns consistently identify the Hindu Kush as the transition between Palearctic and Oriental species, and as a localized hotspot of endemism (Fig. 1). Palearctic taxa reach their southeastern range limits along the massif (*Pelophylax, Bufotes*), whereas Oriental taxa do not extend beyond its southern foothills (*Duttaphrynus, Euphlyctis, Hoplobatrachus*) (Figs. 1–2). In addition, the Hindu Kush hosts relict endemic species of both Palearctic (*Paradactylodon*) and Oriental affinities (*Chrysopaa*), underscoring its dual biogeographic legacy.

**Figure 1.**
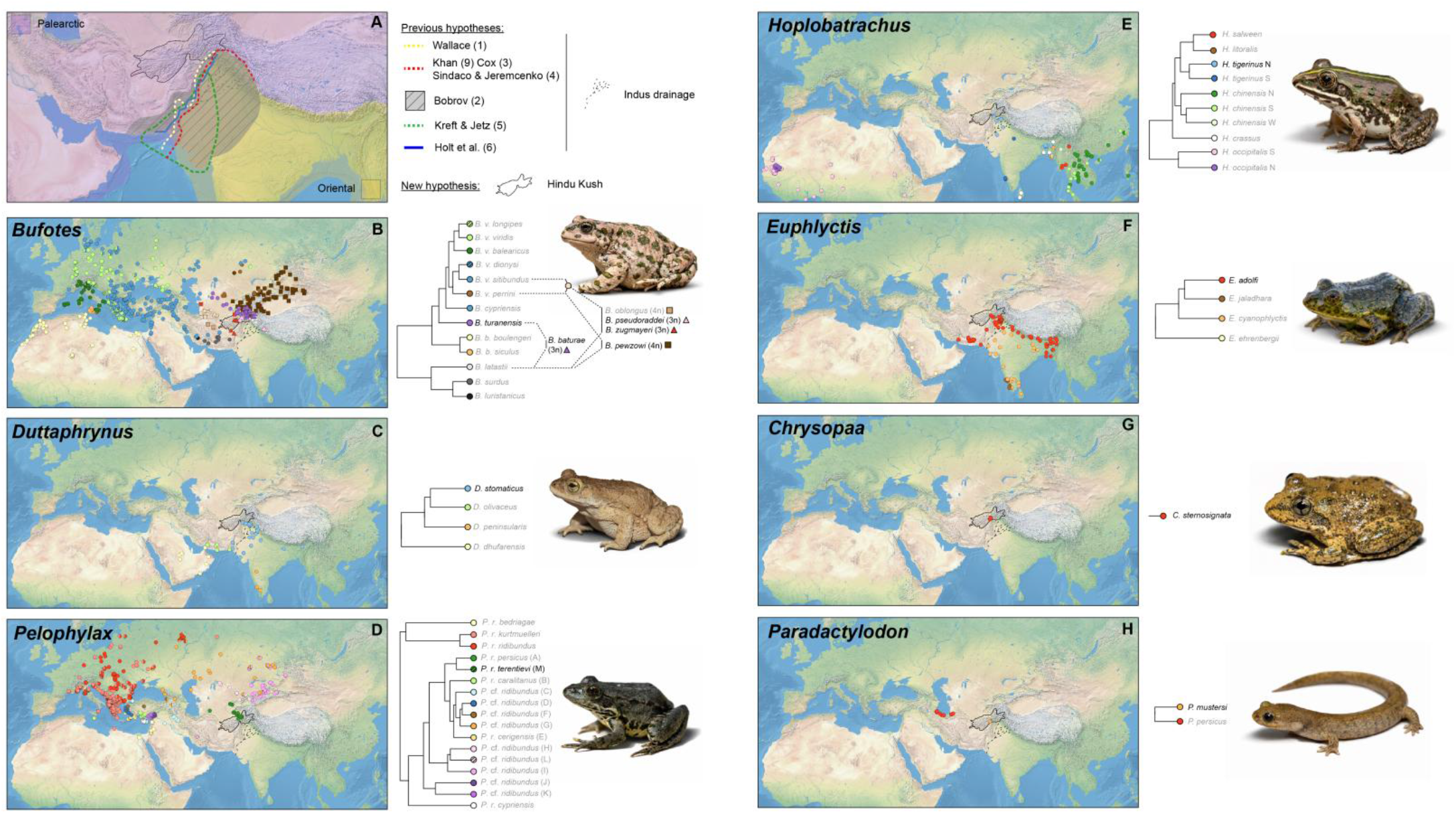
Biogeographic hypotheses for the Palearctic-Oriental western boundary and diversification patterns in the seven amphibian genera occurring in the Hindu Kush region. (A) Hypotheses from literature and this study. (B–H) Phylogeography of amphibian lineages. Maps integrate DNA barcoding of newly sampled Central Asian material with published sequence data. Trees summarize lineage relationships and latest taxonomic assignments; lineages reaching the Hindu Kush are highlighted. In *Bufotes*, hybrid origin of allopolyploid taxa are indicated, with ploidy levels represented by distinct symbols. Photo credit: Daniel Jablonski.

**Figure 2.**
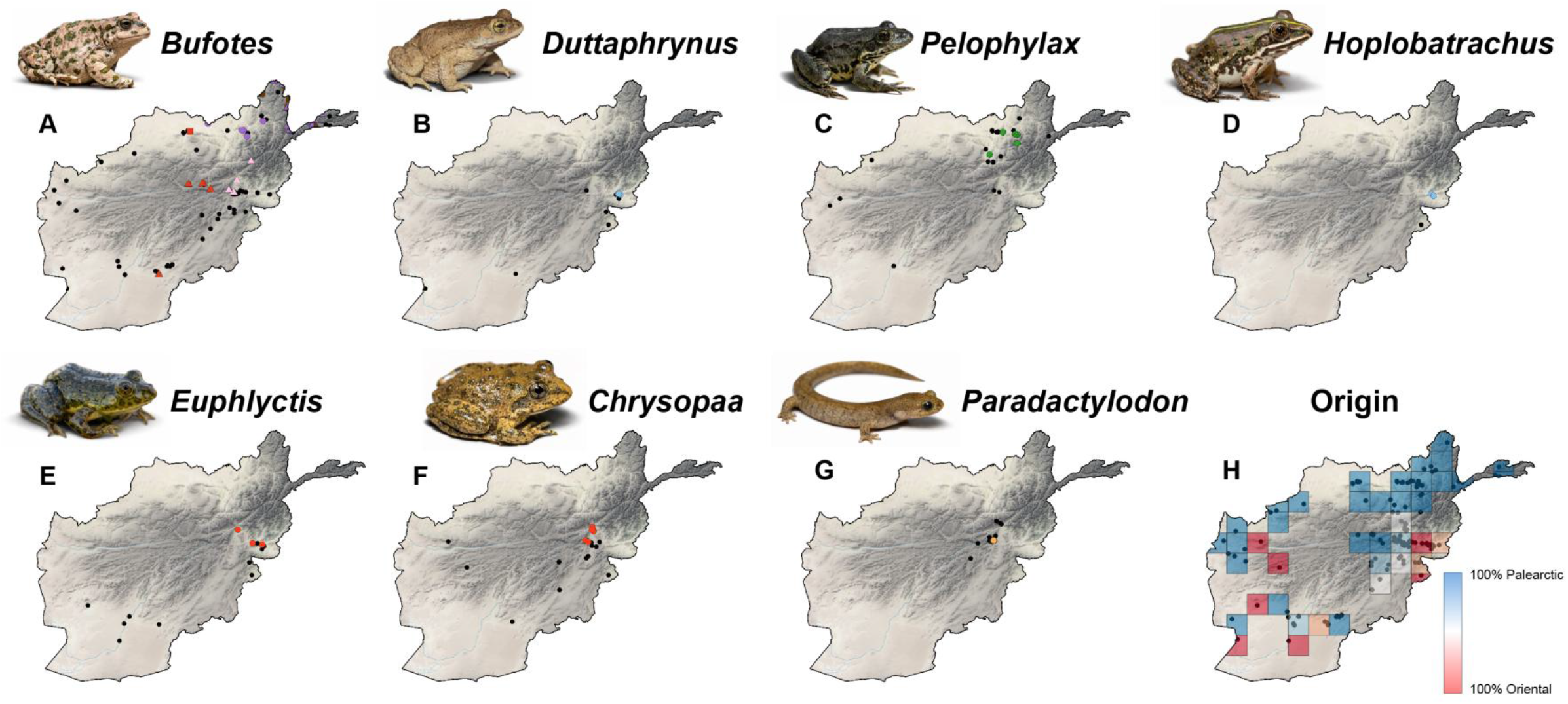
Amphibian updated distributions and biogeographic composition in Afghanistan. (A–G) Occurrence records for each genus. Black dots indicate records lacking genetic data, whereas colored dots represent barcoded samples (see lineage labels in Fig. 1). (H) Proportion of Palearctic vs. Oriental genera within 100×100 km grid cells. Photo credit: Daniel Jablonski.

Overall, three main biogeographic patterns emerge. First, predominantly lowland genera are represented by single, weakly structured lineages that do not cross the Hindu Kush. North of the mountain, the Palearctic water frog *P. ridibundus* occurs in northern Afghanistan as the Amu-Darya subspecies *P. r. terentievi*; scattered (unsampled) western Afghan records (Fig. 2) likely correspond to its sister subspecies *P. r. persicus*, confirmed in adjacent Iran (Fig. 1). South of the massif, the Oriental taxa *D. stomaticus* (Indian Marbled Toad), *E. adolfi* (Northern Skittering Frog), and *H. tigerinus* (Indian Bullfrog) belong to lineages largely associated with the Gangetic plains that reach their westernmost limits in the subtropical lowlands of eastern Afghanistan (Fig. 1). In all cases, distributions terminate against the mountain front, and Oriental lineages cross the Indus River without phylogeographic structure.

Second, *Bufotes* green toads display markedly higher diversity and spatial structuring around the Hindu Kush. Up to five lineages occur, including Central Asian diploid (2n) forms and recently evolved allopolyploid (3n/4n) hybrids (Fig. 1). These are geographically segregated around the massif: 3n *B. zugmayeri* in central-southern areas, 3n *B. pseudoraddei* in the southeast, 3n *B. baturae* and 4n *B. pewzowi* in the east, and 2n *B. turanensis* in the north.

Although *Bufotes* do not extend beyond the southern slopes of the Hindu Kush, two high-elevation species (2n *B. latastii* and 3n *B. pseudoraddei*) marginally penetrate the Oriental realm further east along the Himalayan belt, reaching as far as Zamda County in Chinese Tibet. This mosaic emphasizes the massif as both a barrier and a recent diversification/contact zone between hybridizing lineages.

Third, endemic taxa indicate ancient colonization and long-term persistence within the Hindu Kush. The Karez Frog *C. sternosignata* – the sole representative of its genus – occurs primarily in the southern Hindu Kush of Afghanistan and Pakistan; its closest relatives belong to Himalayan genera of the *Paa* tribe. Conversely, the Afghan Salamander *P. mustersi*, confined to the central Hindu Kush, represents a relict of formerly widespread temperate lineages in Central Asia, with closest relatives in Hyrcania of northern Iran (*P. persicus*) and at the Kazakhstan-China border (*Ranodon sibiricus*). Together, these endemics highlight the massif as both a refugium and a long-term evolutionary reservoir at the interface of the two realms.

More broadly, amphibian occurrences in Afghanistan updated with new field records confirm a Palearctic-Oriental transition centered on the eastern Hindu Kush orography (between Nangarhar and Baghlan Provinces) and its adjacent arid systems (notably the Ragistan Desert) (Fig. 2).

## Discussion

Our data resolve the Palearctic-Oriental interface in Central Asia as a geographically restricted transition aligned with the eastern Hindu Kush and its associated arid systems – not with the Indus River Valley. The concordant north-south partitioning observed across independent amphibian lineages indicates a community-level turnover concentrated along and across the mountain front. Oriental taxa repeatedly reach the southeastern foothills via subtropical corridors but fail to expand across the massif, whereas Palearctic lineages remain largely confined to northern slopes and adjacent basins, or to the massif itself. Even in the complex *Bufotes* system, lineage partitioning clusters around the Hindu Kush and adjacent Western Himalayan ranges rather than with the Indus drainage. Together, these patterns support a spatially coherent boundary rather than the diffuse transition zone hypothesized since Wallace (1).

Such a configuration is consistent with predictions from mountain biogeography. Where steep orography intersects strong hydroclimatic gradients, mountains can simultaneously promote persistence and restrict dispersal, with barrier strength mediated by rain-shadow aridity, thermal regimes, and habitat discontinuity rather than elevation alone (11). Afghanistan exemplifies such filtering: monsoon-influenced forests and productive river corridors are largely confined to the eastern margin, whereas extensive arid basins dominate the interior (10,12). For amphibians, whose distributions are tightly constrained by water availability, this landscape generates inherently asymmetric permeability. In this context, large rivers need not function as persistent realm boundaries; instead, they may facilitate corridor-like dispersal across otherwise inhospitable terrain, as documented for other water-associated vertebrates in Central Asia (13). The absence of a phylogeographic discontinuity across the Indus for the surveyed species – the same lineages occurring east and west of the river – supports this interpretation. At the same time, the presence of relict endemic taxa confined to the Hindu Kush underscores its additional role as a long-term refugium, where lineages of both Palearctic and Oriental affinities have persisted through past climatic oscillations. Because the massif lies within predominantly arid and montane landscapes where amphibian diversity is intrinsically low, its dual role as both a refugium for endemic lineages and a barrier to cross-realm dispersal produces an especially sharp and detectable lineage turnover, making the region an unusually clear natural laboratory for resolving realm boundaries.

By integrating unique genetic sampling from Afghanistan, long underrepresented in regional syntheses despite its key biogeographic position (10,14), our study demonstrates that at diversity margins, inter-realm limits may be structured more consistently by mountain in generally arid environment than by even major river systems, highlighting the disproportionate importance of under-sampled arid-montane regions in refining global biogeographic frameworks.

## Materials and Methods

### DNA barcoding

Amphibian lineages were identified using mitochondrial DNA barcoding based on 16S rRNA and/or, other commonly used markers (ND2, D-loop, COI). Reference datasets were assembled from published sequences and associated metadata, some pre-compiled (15–20), altogether spanning 6,466 individuals (1,873 locality-genus combinations) from 153 studies; *SI Appendix*).

Eighty new DNA tissue samples from five genera (31 locality-genus combinations) were collected across the Hindu Kush in Afghanistan and adjacent Pakistan during field expeditions of DJ conducted between 2018 and 2025. DNA extraction, amplification, and sequencing of the targeted markers followed established protocols (*SI Appendix*). Sequences were aligned manually to the reference datasets and analyzed using maximum-likelihood phylogenetic inference to assess lineage identity and structure. Reference and newly generated material for each group are listed in *SI Appendix*.

### Species occurrence and turnover in Afghanistan

To infer species distributions and turnover within Afghanistan, genetic assignments were combined with validated occurrence records from field surveys, museum vouchers, and literature sources. A total of 179 genus-locality combinations were retained (*SI Appendix*). These data were used to calculate the proportion of Palearctic versus Oriental species within 100×100 km grid cells using QGIS 3.24.3.

## Acknowledgments

We thank the Ministry of Higher Education, Kabul, Afghanistan, and the Ministry of Agriculture, Irrigation and Livestock, Kabul, Afghanistan, for granting permission to conduct field research in the country. We are grateful to Robert Wilson of the Smithsonian Institution, National Museum of Natural History, USA, and Carol Spencer of the Museum of Vertebrate Zoology, Berkeley, USA, for providing tissue samples from their specimen collections. We also thank Hashmat Kamal Fahim, as well as numerous kind individuals, for their assistance with fieldwork, official documentation, and information. We are grateful to Jana Poláková for her work in the DNA laboratory. DJ was supported by the Scientific Grant Agency of the Slovak Republic (VEGA 1/0391/25) and by an EU NextGenerationEU scholarship through the Recovery and Resilience Plan for Slovakia (project no. 09I03-03-V04-00306).

## Author Contributions

D.J. conceived the study and, together with C.D., designed research; all authors performed research; D.J. and C.D. produced or analyzed data; D.J. and C.D. wrote the paper; all authors reviewed and edited the paper.

## Competing Interest Statement

No competing interests.

## Notes

### Competing Interest Statement

The authors have declared no competing interest.

