## Supporting Information for "The Hindu Kush, not the Indus Valley, divides amphibian biogeographic realms"

**Supporting Information Text**

**Extended Methods**

*DNA barcoding details*

Published mitochondrial barcoding data were compiled for markers that differentiate species and major intraspecific lineages of the target amphibian genera (Dataset S1). This involved: (i) harvesting individual and/or haplotype sequences from GenBank and other sources (including publication-associated alignments) (ii) incorporating barcoding information from alternative approaches (e.g., mitotyping without sequencing using lineage-specific restriction enzymes, or information reported for unpublished sequences); and (iii) linking lineage assignments to specimen origins by cross-checking publications and GenBank records. For several groups, we used recently precompiled datasets as starting points. Sequences were handled under SeaView 5 (1).

For *Bufotes*, we updated a 16S rRNA/D-loop dataset (2), now totaling 2,445 individuals from 732 localities, based on 25 studies. Reference alignments comprised individual sequences, namely 1,323 for D-loop (865 bp aligned) and 440 for 16S (449 bp aligned). These sequences distinguish 18 lineages representing most known diploid and allopolyploid taxa, including undescribed polyploid forms.

For *Duttaphrynus*, we updated a 16S dataset (3), now totaling 284 individuals from 157 localities, based on 49 studies. The reference alignment comprised 284 individual sequences (558 bp aligned). These distinguish 23 lineages representing 17 species arranged in two deeply-diverged clades (*Duttaphrynus* and *Firouzophrynus*). Barcoding results focused on the *Firouzophrynus* clade, represented in the study area.

For *Pelophylax*, we considered an ND2 dataset (4) restricted to Western Palearctic lineages, totaling 3,136 individuals from 678 localities, based on 26 studies. The reference alignment comprised 444 haplotype sequences (1,038 bp aligned). These distinguish all Western Palearctic taxa and their many intraspecific lineages. Barcoding results focused on the widespread *P. ridibundus*, represented in the study area.

For *Hoplobatrachus*, we built a 16S dataset de novo, totaling 310 individuals from 178 localities, based on 45 studies. The reference alignment comprised 310 individual sequences (523 bp aligned). These distinguished all species of the genus, as well as deeply diverged lineages within several of them.

For *Euphlyctis*, we updated a 16S dataset (5), now totaling 253 individuals from 110 localities, based on 30 studies. The reference alignment comprised 60 haplotype sequences and 7 individual sequences (481 bp). These distinguish 10 species-level lineages arranged in two deeply-diverged clades (*Euphlyctis* and *Phrynoderma*). Barcoding results focused on the *Euphlyctis* clade, represented in the study area.

For *Chrysopaa*, we reviewed compiled 16S and COI sequences (6), representing eight individuals from five localities, based on two studies.

For *Paradactylodon*, we reviewed compiled COI sequences (7), totaling 30 individuals from 13 localities, based on two studies.

New sequences were generated from Hindu Kush specimens for *Bufotes* ($n$ = 63 from 18 localities), *Duttaphrynus* ($n$ = 3 from three localities), *Pelophylax* ($n$ = 5 from five localities), *Hoplobatrachus* ($n$ = 2 from two localities) and *Euphlyctis* ($n$ = 7 from three localities) (Dataset S1). DNA was isolated with the E.Z.N.A.® Tissue DNA Kit. PCR were performed as follows. For 16S (2,8), for D-loop (2) and for ND2 (9).

Sequences were aligned manually to the reference alignments, available in Dataset S2. Phylogenetic positions were inferred from maximum-likelihood tree reconstructions with IQ-TREE 2 under default settings (10), as provided in Dataset S3. Barcoding data were mapped under QGIS 3.24.3.

For visualization, evolutionary relationships among focal species/lineages (Fig. 1) were adapted from the trees obtained by published studies, based on additional (notably phylogenomic) markers (2–7, 11).

*Occurrence records*

Validated amphibian records in Afghanistan (179 genus-locality combinations) are provided in Dataset S4. Records were mapped alongside barcoding data under QGIS 3.24.3. To calculate the proportion of Palearctic versus Oriental species, we (i) generated a 100×100 km grid with the QGIS function *create grid*; (ii) counted the number of different genera present in each grid cell with the function *Join attributes per location (summary)*); and (iii) quantified the proportion of those of Oriental origin using the *Field Calculator*.

**Datasets: can be made available upon demand.**

**Dataset S1**
Barcoding data for each genus.

**Dataset S2**
Reference alignments and new sequences.

**Dataset S3**
Raw tree files of each IQ-TREE analysis

**Dataset S4**
Updated amphibian records in Afghanistan
